# AI-Enabled Classification of Head and Neck Tumors using Convolutional Neural Networks with Dynamic Optical Contrast Imaging

**DOI:** 10.1101/2022.10.26.513814

**Authors:** Ren Odion, Laith Mukdad, Yazeed Alhiyari, Kenric Tam, Ramesh Shori, Tuan Vo-Dinh, Maie A. St. John

## Abstract

**Background:** Recent advances in artificial intelligence (AI) in the field of imaging have resulted in new opportunities for automated tumor detection and margin assessment. In particular, AI deep learning techniques such as the Convolutional Neural Network (CNN) have greatly advanced the field of computer vision. Here we introduce the application of a CNN model for use with Dynamic Optical Contrast Imaging (DOCI), an imaging technique developed by our group that creates a unique molecular signature on tissue targets by obtaining the autofluorescence decay of several spectral bands in the UV-Vis range.

**Methods:** 21 patients undergoing surgical resection for tonsillar squamous cell carcinoma (SCC) were identified on a prospective basis. DOCI images were analyzed and compared to the pathology results as ground truth. A CNN model was used to segment sections of DOCI images and provide a percentage chance of tumor presence, allowing for automated tumor margin delineation without a-priori knowledge of the tumor tissue composition.

**Results:** CNN outputs yielded a 99.98% confidence in classifying non-tumor tissue and 76.02% confidence in classifying tumor tissue.

**Conclusions:** Our results indicate that a CNN-based classification model for DOCI allows for real-time analysis of tissue, providing improved sensitivity and accuracy of determining true margins and thus enabling the head and neck cancer surgeon to save healthy tissue and improve patient outcomes.

## INTRODUCTION

Over 450,000 patients are diagnosed with head and neck squamous cell carcinoma (HNSCC) every year.^1^ Prognosis of HNSCC patients with depends heavily on complete tumor resection. Positive margin status, commonly defined as any viable tumor at the edge of the resected tumor, is associated with significantly decreased survival due to local or regionally persistent or recurrent disease.^2,3^ The overall number of head and neck cancer patients with true positive tumor margins after cancer surgery can range from 10-25%.^4^

Intraoperatively, surgeons determine gross tumor margins through palpation and visual inspection. After a tumor is presumed to be removed in its entirety, the surrounding tissues are sampled by frozen section in an effort to ensure that no microscopic disease is left behind. An assessment of microscopic margins is performed using frozen section (FS) histologic pathology, a process that can take 30 minutes to 1 hour. The efficacy of this approach varies widely and is subject to sampling error. Multiple studies have demonstrated that FS utilization does not improve survival or locoregional control and furthermore, is not predictive of final margin status.^3-5^ The methodology by which FS are collected—whether tumor-bed driven or specimen driven—can also alter margin outcome.^6^ Thus, improving intraoperative detection of tumor margins will be key to optimizing treatment and outcomes.

Our group has developed and previously described Dynamic Optical Contrast Imaging (DOCI), a novel imaging modality that acquires temporally-dependent measurements of tissue autofluorescence.^5-9^ Additionally, in fresh ex-vivo experiments, our group showcased that the DOCI system reliably differentiates HNSCC from surrounding normal tissue and facilitates specific tumor localization [**Figure 1**].^5^ DOCI images are captured in real-time, do not require administration of contrast agents, and offer an operatively wide field of view. DOCI probes the intrinsic contrast of fluorescence lifetime of different endogenous fluorophores within tissue, creating a unique molecular map that can be used to determine cancer margins. This margin delineation process must then be cross-referenced with histology slides serving as the ground truth. Previous work has been done to statistically quantify DOCI images for tissue classification ranging from basic logistic regression to artificial neural networks (ANNs).^10^

**Figure 1:**
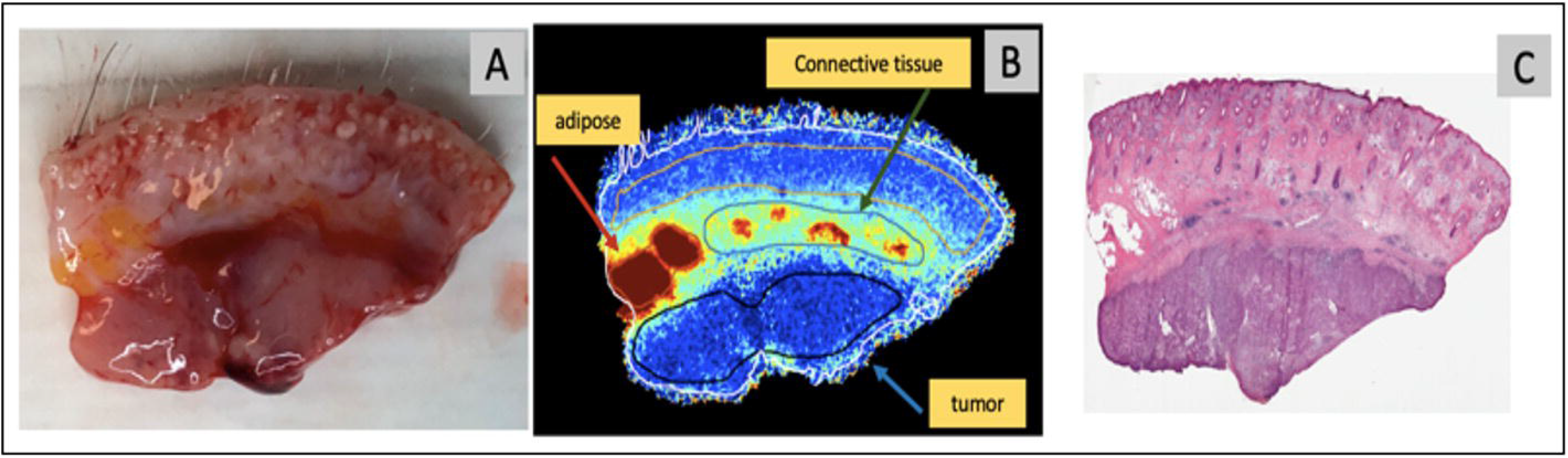
DOCI can differentiate cancer from muscle and surrounding non-cancer tissues. In figure 1, we see a specimen that contains part normal tissue and part cancer. A) RGB image of grossly resected specimen. B) DOCI image of specimen. C) H&E staining of specimen. We see that even within one specimen, DOCI can readily discriminate between normal tissue and cancer tissue. Histology and DOCI are concordant.

Deep learning has gained wide usage in image classification in a wide area of fields from object detection to image classification and segmentation.^11^ Such multi-layered machine learning networks work by extracting classifying features from inputs such as RGB images. Convolutional neural networks (CNNs) are a particularly popular neural network in which convolutional layers act as feature extractors and pooling layers subsequently down sample those feature maps, leading to network sizes that are relatively small and computationally feasible compared to traditional fully connected ANNs.^11^ Such feature extraction and classification make CNNs an ideal model for image classification for tumors in particular. Recently, deep learning has also been used with input data from hyperspectral images wherein the image are obtained from different spectral regions other than the usual visible range RGB.^12,13^ We have also developed CNN models for spectral data from plasmonic nanoprobes using Surface-Enhanced Raman Spectroscopy in quantifying HNSCC from extracted samples.^14^ In this study, we enhance the performance of the DOCI system by developing a similar CNN model trained on spectrally separated DOCI images of tonsillar SCC tissue. CNN models trained on DOCI images containing molecular information serve as additional features for better tumor classification that deep learning will leverage. Using this CNN model, we can create an automated image processing pipeline wherein DOCI images are classified based on the likelihood of tumor presence on the image.

## METHODS

This study was approved by the institutional review board of the University of California Los Angeles (UCLA). Patients undergoing surgical resection for tonsillar squamous cell carcinoma (SCC) were identified on a prospective basis. Before surgery, all patients signed a written consent form for involvement in the study. Twenty-one patients were included. All specimens were confirmed to be squamous cell carcinoma on final pathology. After surgical resection, specimens were immediately sectioned into multiple fresh samples containing tumor and contiguous normal tissue of suspect lesions, and the tissue was transferred by courier to the UCLA Translational Pathology Core Laboratory where the DOCI system was housed, and the specimens were immediately imaged. DOCI imaging was performed within 20 minutes, and imaging acquisition required approximately 90 seconds.

### Dynamic optical contrast imaging instrumentation

The instrumentation and setup for the DOCI system have been previously published.^15^ The DOCI technique utilizes a 365 nm UV LED as an excitation source to produce contrast between fluorophores of different decay rates. Fundamentally, DOCI relies on the fact that the longer lifetime fluorophores produce more signal than the shorter lifetime fluorophores when referenced to their steady state fluorescence. To further assist with differentiating the subtle autofluorescence signal indicative of different tissue types, the autofluorescence signal is captured using different spectral filters (bands/channels) [**Figure 2**].

**Figure 2:**
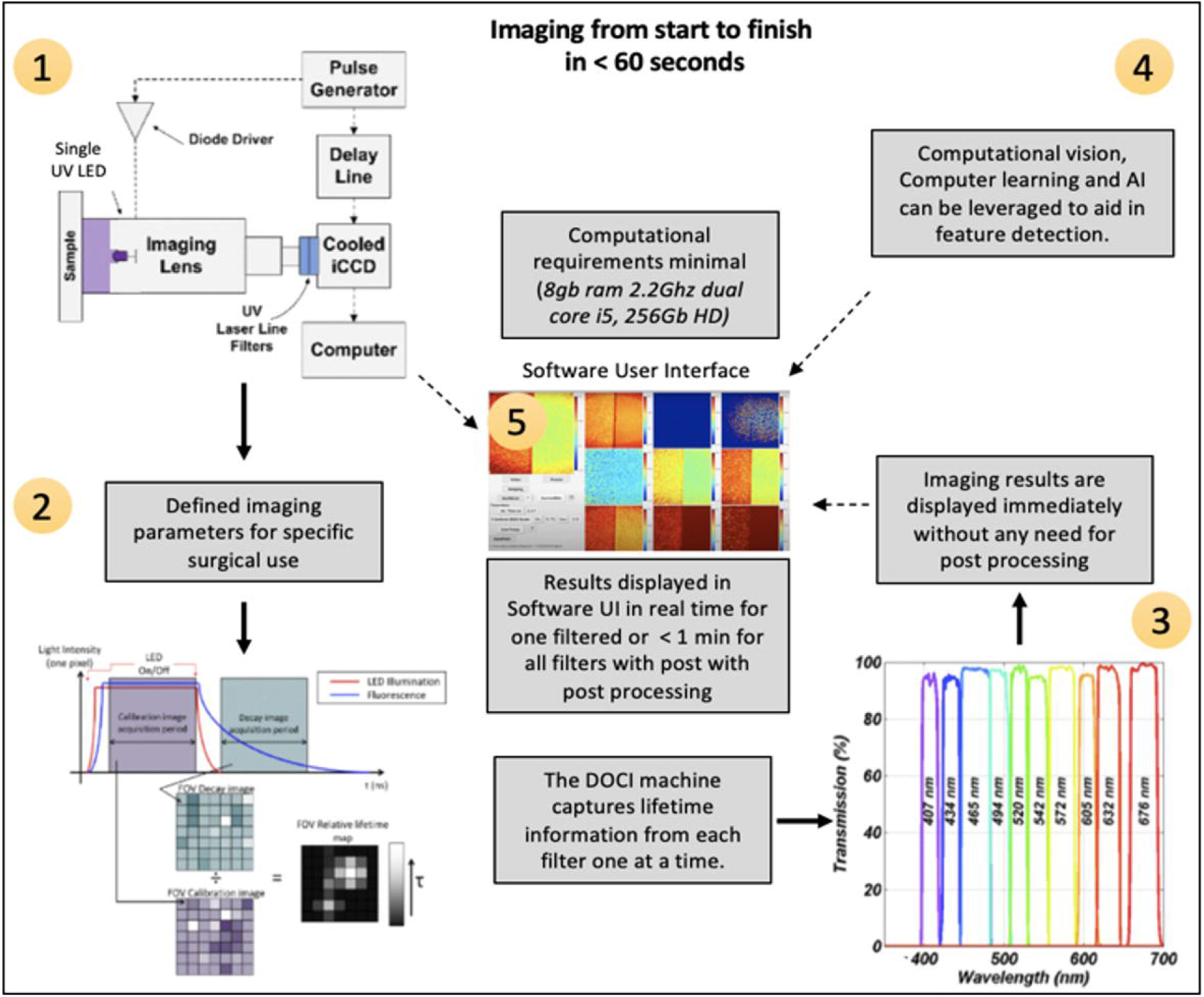
DOCI Concept and Process. 1) Block diagram of DOCI; 2) Image generation algorithm starts with defined imaging parameters for specific surgical use; 3) Images are captured at the associated wavelengths; 4) AI and Computer aided feature detection can be added and in the software UI; 5) The software UI is easy to use (need only 3 clicks for image or video acquisition. Imaging results are displayed immediately without any need for post processing.

### DOCI image processing and statistical analysis

DOCI image processing and calculation has been detailed previously by our group.^5^ In brief, a DOCI image was calculated for 9 filter channels, and an intensity image is computed for 9 BPF channels. An aggregate intensity measurement was recorded from the moment the illumination started to decrease until the fluorescence ceased. A DOCI image was calculated by normalizing the aggregate fluorescence decay intensity by the aggregate stead-state fluorescence intensity.

Subsequently, a head and neck surgeon and the head and neck pathologist were presented with both the registered visible image and the whole-section histology displayed on a computer screen. Both the surgeon and the pathologist were blinded to the DOCI images. Features from the visible images were compared with features in histology, and closed contours were manually drawn on the visible images representing tumor and non-tumor as indicated in **Figure 3**.

**Figure 3:**
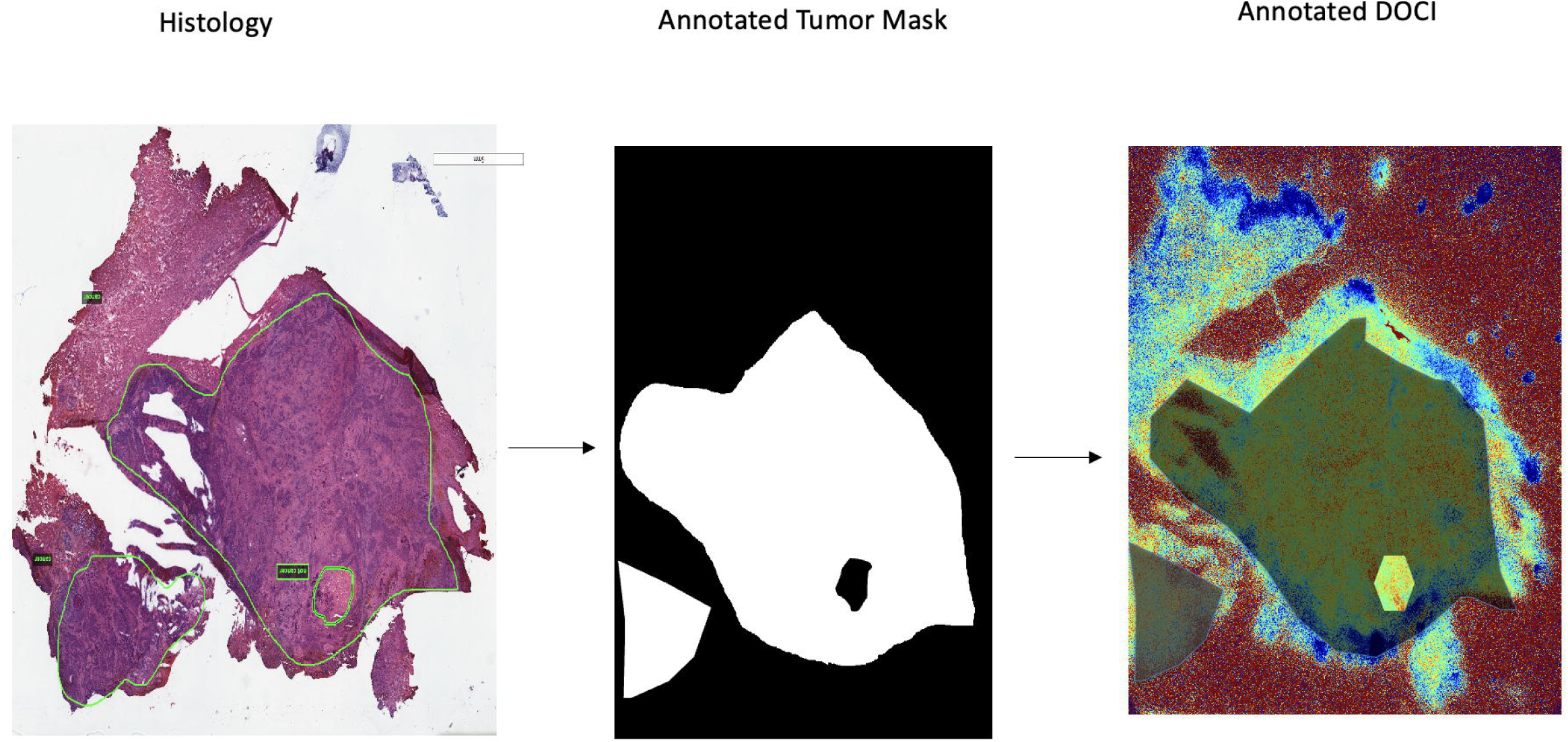
Workflow for tumor annotation pipeline. Histology slides are initially annotated and used as a reference to create a tumor mask used in DOCI image classification (shown as darkened regions on the DOCI image).

For each tissue sample, a region of interest (ROI) corresponding to a tissue type identified on frozen histology was selected. The average DOCI value for each ROI was subsequently calculated. DOCI values for each tissue type were normalized against the tumor DOCI value. After confirming a normal distribution, a one-sample t-test was utilized for statistical analysis between different tissue types and cancer. Data analysis was performed using MATLAB R2021b and Microsoft Office Excel. The DOCI annotation process is visually summarized in **Figure 3**.

### Convolutional Neural Network

The CNN network was developed using the TensorFlow framework in Python. The model consists of 3 convolutional layers for feature extraction. A fully connected dense layer and sigmoidal output layer is set to create a binary output inference. Several optimizations such as the inclusion of a 10% dropout rate just before the dense layer allows for a reduction of overfitting. Finally, the Adam optimizer with a mean squared error loss was used for the prediction of tissue classification. A simplified visualization of the CNN model is shown in **Figure 4**.

**Figure 4:**
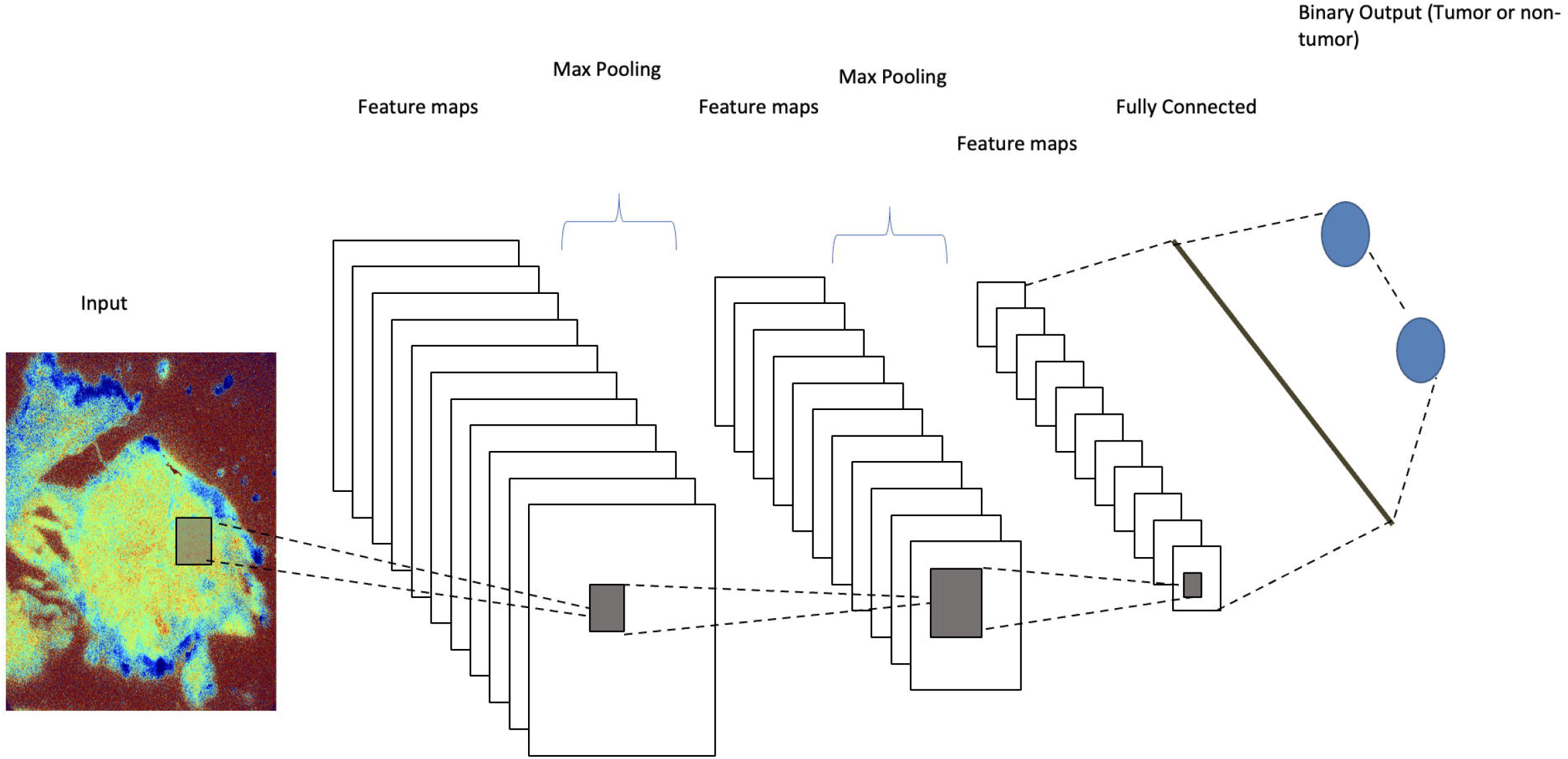
CNN network with DOCI image input and binary tumor classification output

### Creation of Training and Validation Sets

DOCI images of tonsillar SCC tissue were used as the input for the CNN model. DOCI images consist of spectrally separated images from 400 nm to 800 nm. A total of X unique samples was collected. Each sample contains 20 DOCI tumor annotated images, and each DOCI image is an 8-bit RGB image that is 900×900 pixels in resolution. Each DOCI image is split to 9 equal parts of 300×300 pixel resolution and binned to either tumor or non-tumor based on the tumor annotation masks associated with each image. Notably if more than 5% of the pixels in the image segment is tumor annotated, such images are classified as tumors. A total of 683 tumor and 2800 non-tumor images were used with a 5 to 1 training to validation dataset split. All image processing were conducted using Python with the Sci-Kit Image library. **Figure 5** shows the image processing and data segregation pipeline with DOCI images.

**Figure 5:**
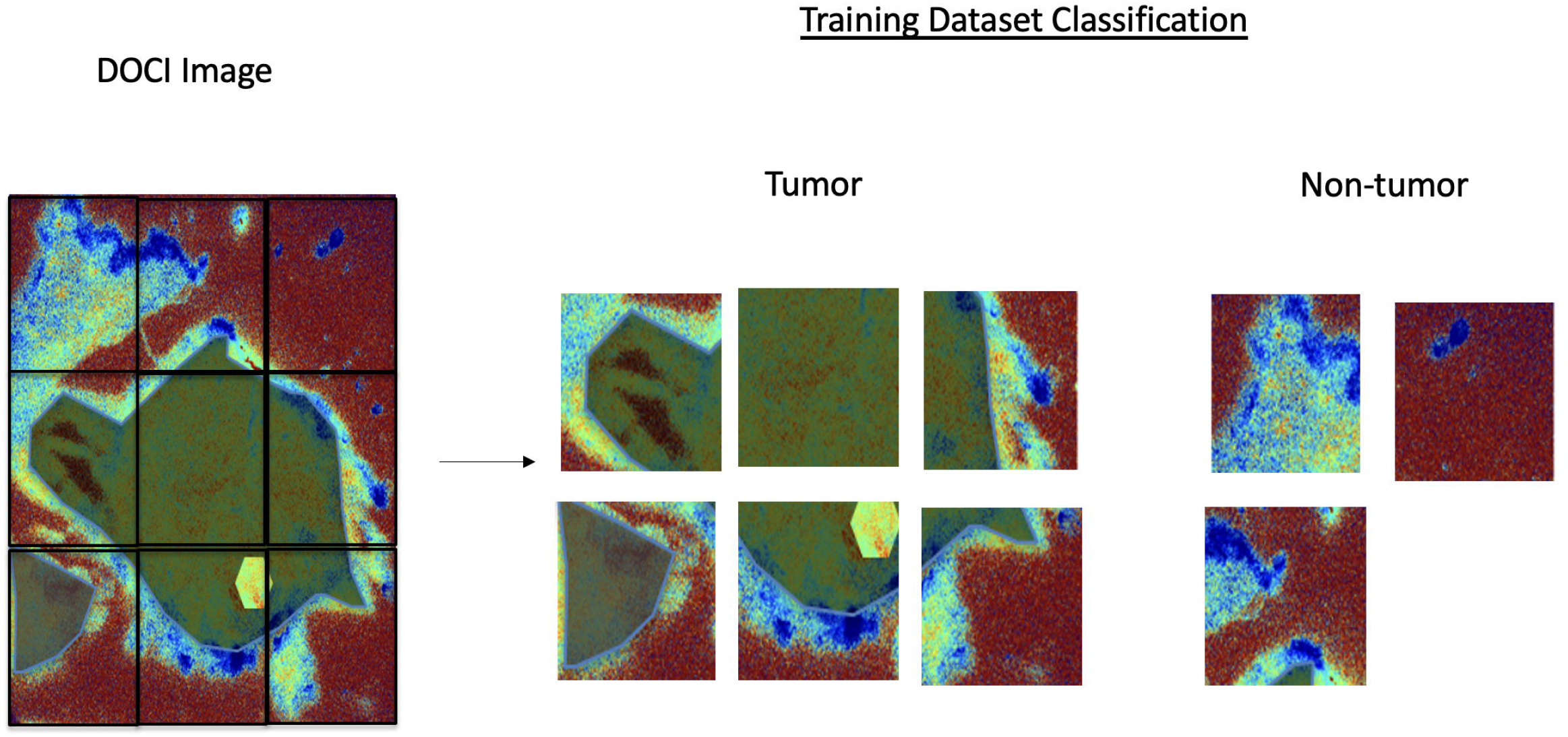
Dataset creation using split subsections of DOCI image. Each subsection is classified as tumor if more than 5% of the pixel map comprises of the tumor annotated mask.

### CNN Training

The CNN model was trained using a local workstation housing two RTX 2060s (Nvidia, Santa Clara, US). A training time of 10 minutes was achieved with 50 epochs. The resulting training performance metrics were plotted using Python with the Matplotlib graphing library.

## RESULTS

### DOCI Lifetime Mapping

DOCI lifetime mapping produced statistically significant differences in contrast between tumor and non-tumor across most multiple wavelengths. A decrease in fluorescence lifetime was observed in malignant tissue that is consistent with the short lifetimes reported for biochemical markers of tumors (**Figure 6**). Statistical significance by Wilcoxon rank-sum test (P < .05) between tumor and non-tumor.

**Figure 6:**
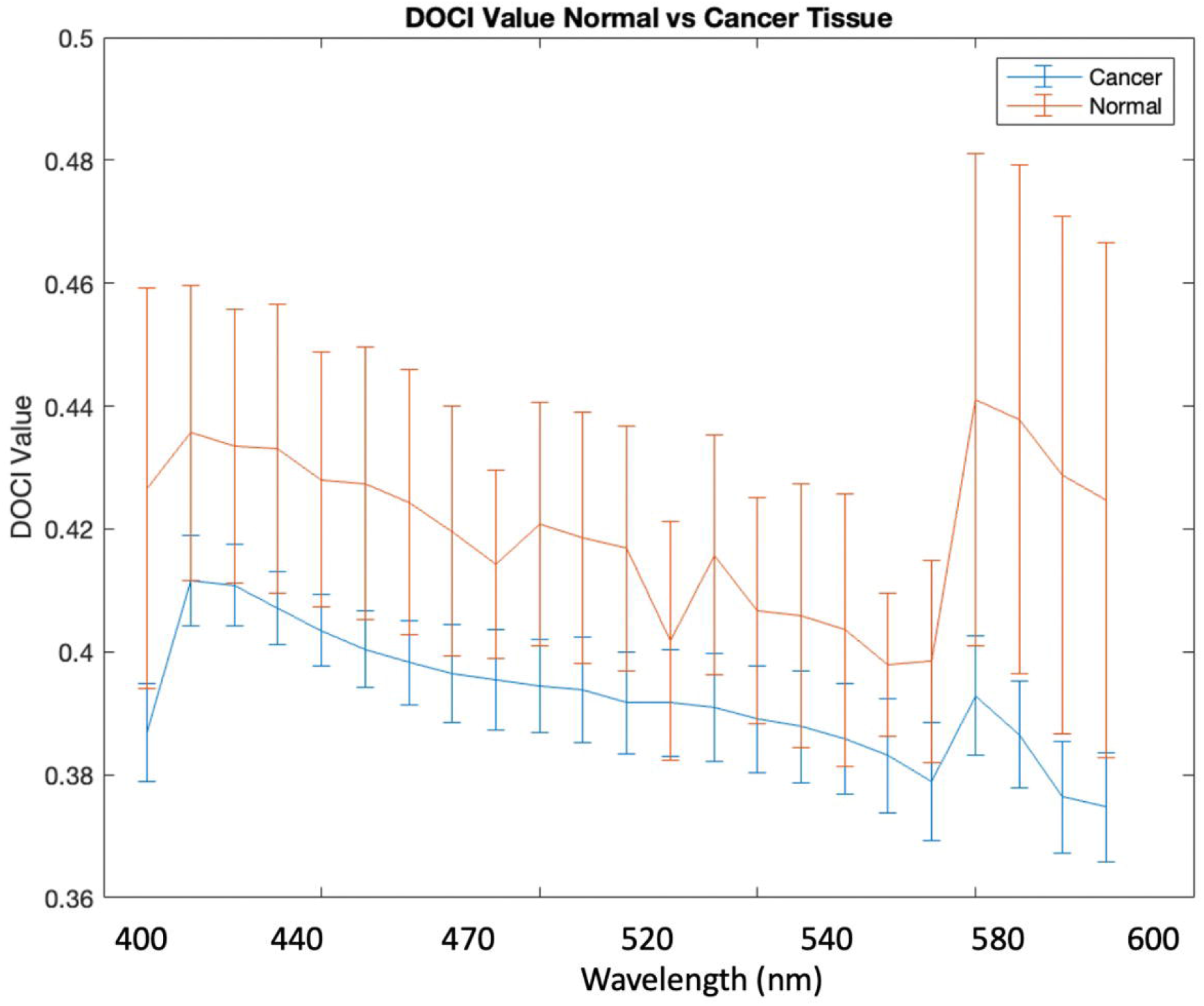
Average relative lifetimes of cancer and normal tissue as a function of wavelength.

### CNN Training and Evaluation

After collection of DOCI images, a python script is used to automatically label and distribute images as tumor or nontumor. This training set is used to train the CNN model described in **Figure 4** for 50 epochs. The resulting model performance metrics is displayed in **Figure 7**.

**Figure 7:**
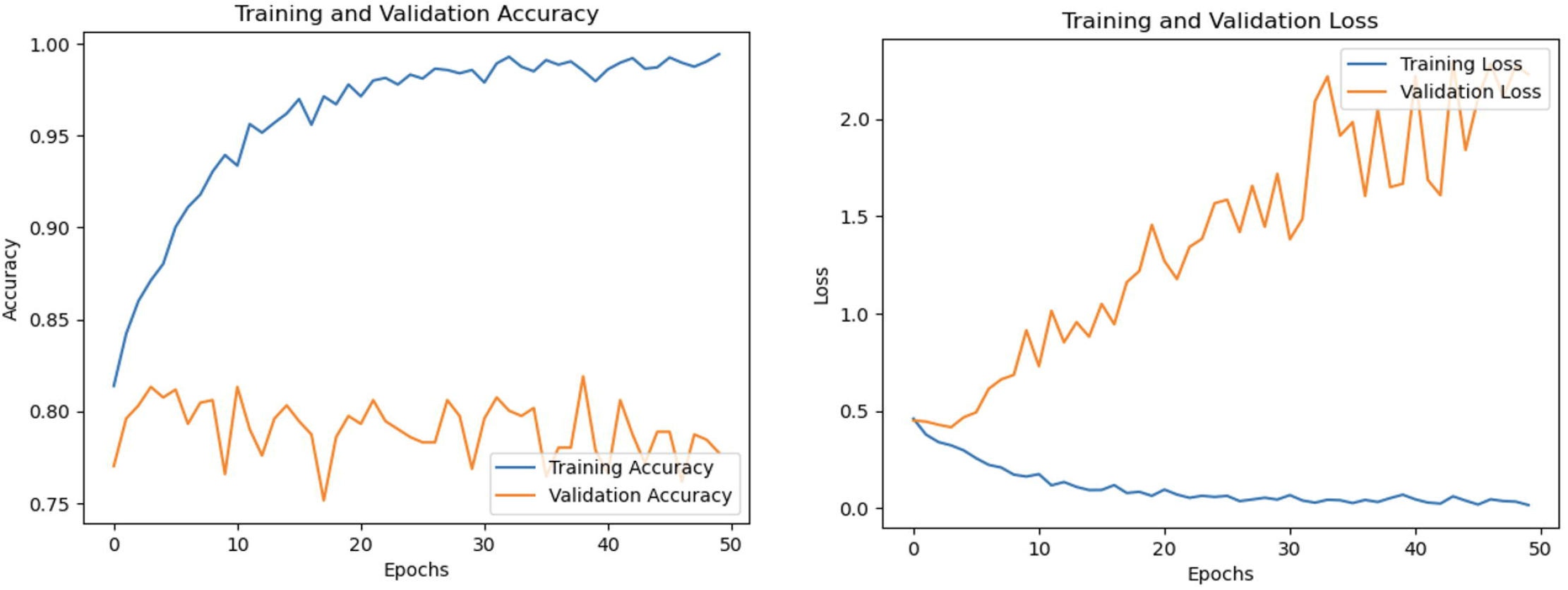
CNN training performance across 50 epochs.

### CNN Inference for Tumor Classification

Inference is demonstrated in **Figure 8**, where an input DOCI image at 405-410 nm is fed into the model. The resulting output are the segmented and classified images with a certain percentage of tumor or non-tumor classification. In **Figure 8**, we see that the section with no tumor annotation is classified as non-tumor with a 99.98% confidence while the section with the tumor is classified as tumor with 76.02% confidence.

**Figure 8:**
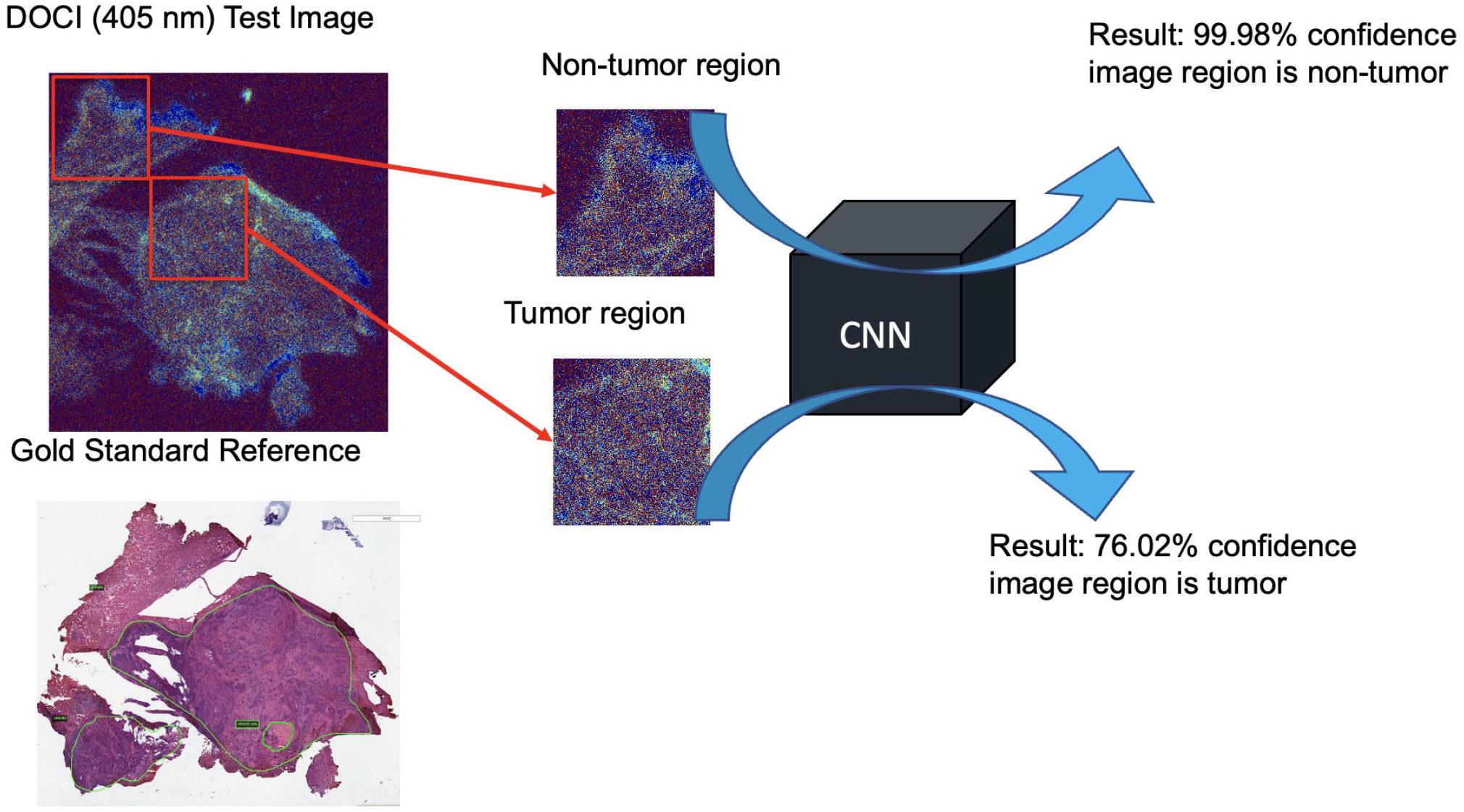
CNN inference on input DOCI image for subsection classification as tumor or non-tumor

## DISCUSSION

The recent momentous drive to apply advanced AI technologies to diagnostic medicine has the potential to revolutionize cancer surgical care by allowing it to be more precise and personalized. CNNs are one of the most popular AI algorithms used for deep learning. While CNNs have progressed the field of oncologic pathology, their use with respect to head and neck oncology remains in the nascent stage. Moreover, although AI-based methods have been used to analyze large data registries of head and neck cancer patients, the extent of data analyzed has mostly been limited to demographic, clinicopathologic, or genomic data.^16-19^

This prospective study of 21 patients with tonsil SCC is the first to combine the technologies of DOCI and CNN-based classification models, yielding promising results that further DOCI’s clinical translatability, aiding surgeons with precise head and neck cancer surgical resection. As has already been demonstrated by our previous studies and in the current one, DOCI reliably differentiates head and neck cancer tissue from surrounding normal tissue on the basis of fluorescence decay information.^5-10^ The performance of the DOCI system is enhanced by the CNN-based classification model for automated classification of tumor and non-tumor tissue of DOCI-acquired images. As the CNN model continues to acquire further DOCI-acquired image samples, we anticipate that the tumor region confidence will continue to improve above the current 76.02% confidence mark. Comparably, Choudhry et. al noted a 76% accuracy in delineating breast cancer with an input of 162 breast cancer histological slides, and Mohsen et al noted an 87.2% accuracy in delineating brain cancer tissue with an input of 66 MRI images.^20,21^ It is well established that the accuracy of CNN models improve as larger training data are inputted,^22^ and we anticipate an increased accuracy as we continue to enroll patients into our study. Currently, the CNN model consists of only 3 convolutional layers for extracting features used for the final output tumor classification, resulting in a limited performance on unseen data input as shown in the validation curves of **Figure 7**. However, future studies can leverage more complex CNN architectures such as those with many convolutional layers as VGG16 or ResNet which may have the ability to create a better classification model for identifying tonsillar SCC tissue from non-tumor tissue.

Our works builds upon prior CNN models used for spectral data from plasmonic nanoprobes using Surface-Enhanced Raman Spectroscopy in quantifying head and neck cancer from extracted samples.^14^ In this study, our group augments the performance of the DOCI system by developing a similar CNN model trained on spectrally separated DOCI images of tonsillar SCC tissue. CNN models trained on DOCI images containing molecular information serve as additional features for better tumor classification that deep learning will leverage. Using this CNN model, we can create an automated image processing pipeline wherein DOCI images are classified based on the likelihood of tumor presence on the image. Following this pilot trial, we aim to validate the proposed method in more datasets, when available. Tongue SCC tissue were also collected as part of the study and incorporating this data into the CNN design and training will allow for tumor tissue type classification rather than just tumor or non-tumor.

In this proof-of-concept study, we demonstrate that a CNN-based classification model with DOCI image input can accurately delineate cancer and non-cancer tissue. As a first step in clinical translatability, our CNN model may be used to instantaneously analyze DOCI images that are acquired immediately after tumor resection but before FS biopsies to direct biopsies. Currently, FS margin biopsies are determined through a combination of surgeon experience, visual cues, and palpation. The lack of uniform practice causes geographical misses and some positive margins may not be identified.^3-5^ Our AI-supplemented DOCI imaging technique may be key to achieving negative margins intraoperatively, optimizing oncologic outcomes and improving patient overall survival while preserving healthy tissue and decreasing morbidity.

## CONCLUSION

In this study, we developed a CNN-based classification model for automated classification of tumor and non-tumor tissue in DOCI images to allow for precision surgery. CNN was used to segment sections of a DOCI image and give a percent chance of tumor presence, allowing for automated tumor margin delineation without a-priori knowledge of the tumor tissue composition after *in-vitro* histology analysis. The results indicate that a CNN-based classification model for DOCI allows for real-time analysis of tumor tissue, providing potential to significantly improve the sensitivity and accuracy of determining true margins and thus enabling the head and neck cancer surgeon to save healthy tissue and improve patient outcomes.

## ACKNOWLEDGEMENTS

This study is supported by the US National Institute of Health (grants R01EB028078 and R01 CA220663-06) and the American Academy of Otolaryngology-American Head & Neck Society Alando J. Ballantyne Resident Research Pilot Grant.

## REFERENCES

1. Mignogna MD, Fedele S, Lo Russo L. The World Cancer Report and the burden of oral cancer. Eur J Cancer Prev. 2004;13(2):139–42. Epub 2004/04/22. doi: 10.1097/00008469-200404000-00008. PubMed PMID: 15100581.

2. Shapiro M, Salama A. Margin Analysis: Squamous Cell Carcinoma of the Oral Cavity. Oral Maxillofac Surg Clin North Am. 2017;29(3):259–67. Epub 2017/07/16. doi: 10.1016/j.coms.2017.03.003. PubMed PMID: 28709529.

3. Na HS, Kwon HK, Shin SC, Cheon YI, Seo M, Lee JC, Sung ES, Lee M, Kim IJ, Kim BH, Lee BJ. Clinical outcomes of T4a papillary thyroid cancer with recurrent laryngeal nerve involvement: a retrospective analysis. Sci Rep. 2021;11(1):6707. Epub 2021/03/25. doi: 10.1038/s41598-021-86226-x. PubMed PMID: 33758286; PMCID: PMC7988054.

4. Zanoni DK, Migliacci JC, Xu B, Katabi N, Montero PH, Ganly I, Shah JP, Wong RJ, Ghossein RA, Patel SG. A Proposal to Redefine Close Surgical Margins in Squamous Cell Carcinoma of the Oral Tongue. JAMA Otolaryngol Head Neck Surg. 2017;143(6):555–60. Epub 2017/03/10. doi: 10.1001/jamaoto.2016.4238. PubMed PMID: 28278337; PMCID: PMC5473778.

5. Tajudeen BA, Taylor ZD, Garritano J, Cheng H, Pearigen A, Sherman AJ, Palma-Diaz F, Mishra P, Bhargava S, Pesce J, Kim I, Sebastian C, Razfar A, Papour A, Stafsudd O, Grundfest W, St John M. Dynamic optical contrast imaging as a novel modality for rapidly distinguishing head and neck squamous cell carcinoma from surrounding normal tissue. Cancer. 2017 Mar 1;123(5):879–886. doi: 10.1002/cncr.30338. Epub 2016 Oct 20. PMID: 27763689.

6. Kim IA, Taylor ZD, Cheng H, Sebastian C, Maccabi A, Garritano J, Tajudeen B, Razfar A, Palma Diaz F, Yeh M, Stafsudd O, Grundfest W, St John M. Dynamic Optical Contrast Imaging. Otolaryngol Head Neck Surg. 2017 Mar;156(3):480–483. doi: 10.1177/0194599816686294. Epub 2017 Jan 24. PMID: 28116982.

7. Pellionisz PA, Badran KW, Grundfest WS, St John MA. Detection of surgical margins in oral cavity cancer: the role of dynamic optical contrast imaging. Curr Opin Otolaryngol Head Neck Surg. 2018 Apr;26(2):102–107. doi: 10.1097/MOO.0000000000000444. PMID: 29517537; PMCID: PMC5846197.

8. Hu Y, Han AY, Huang S, Pellionisz P, Alhiyari Y, Krane JF, Shori R, Stafsudd O, St John MA. A Tool to Locate Parathyroid Glands Using Dynamic Optical Contrast Imaging. Laryngoscope. 2021 Oct;131(10):2391–2397. doi: 10.1002/lary.29633. Epub 2021 May 27. PMID: 34043240.

9. Tam, K., Alhiyari, Y., Huang, S. et al. Label-free, real-time detection of perineural invasion and cancer margins in a murine model of head and neck cancer surgery. Sci Rep 12, 12871 (2022). https://doi.org/10.1038/s41598-022-16975-w

10. Huang, S., Alhiyari, Y., Hu, Y., Tam, K., Han, A. Y., Krane, J. F., Shori, R., St. John, M. A. & Stafsudd, O. Ex vivo hypercellular parathyroid gland differentiation using dynamic optical contrast imaging (DOCI). Biomed. Opt. Express 13, 549 (2022).

11. Sultan, A. S., Elgharib, M. A., Tavares, T., Jessri, M. & Basile, J. R. The use of artificial intelligence, machine learning and deep learning in oncologic histopathology. J. Oral Pathol. Med. 49, 849–856 (2020).

12. Chen, Y., Lin, Z., Zhao, X., Wang, G. & Gu, Y. Deep Learning-Based Classification of Hyperspectral Data. IEEE J. Sel. Top. Appl. Earth Obs. Remote Sens. 7, 2094–2107 (2014).

13. Chen, Y., Jiang, H., Li, C., Jia, X. & Ghamisi, P. Deep Feature Extraction and Classification of Hyperspectral Images Based on Convolutional Neural Networks. IEEE Trans. Geosci. Remote Sens. 54, 6232–6251 (2016).

14. Li JQ, Dukes PV, Lee W, Sarkis M, Vo□Dinh T. Machine learning using convolutional neural networks for SERS analysis of biomarkers in medical diagnostics. Journal of Raman Spectroscopy. 2022 Sep 12.

15. Hu, Y. et al. Design and validation of an intraoperative autofluorescence lifetime imaging device. Imaging Therap. Adv. Technol. Head Neck Surg. Otolaryngol. 11213, 23.https://doi.org/10.1117/12.2560000 (2020)

16. Karadaghy OA, Shew M, New J, Bur AM. Development and assessment of a machine learning model to help predict survival among patients with oral squamous cell carcinoma. JAMA Otolaryngol Head Neck Surg. 2019;145(12):1115

17. Chang SW, Abdul-Kareem S, Merican AF, Zain RB. Oral cancer prognosis based on clinicopathologic and genomic markers using a hybrid of feature selection and machine learning methods. BMC Bioinformatics. 2013;14:170.

18. Patil S, Habib Awan K, Arakeri G, et al. Machine learning and its potential applications to the genomic study of head and neck cancer-A systematic review. J Oral Pathol Med. 2019;48(9):773–779.

19. Li S, Chen X, Liu X, et al. Complex integrated analysis of lncRNAs-miRNAs-mRNAs in oral squamous cell carcinoma. Oral Oncol. 2017;73:1–9

20. Choudhury A, Perumalla S. Detecting breast cancer using artificial intelligence: Convolutional neural network. Technol Health Care. 2021;29(1):33–43. doi: 10.3233/THC-202226. PMID: 32444590.

21. Heba Mohsen, El-Sayed A. El-Dahshan, El-Sayed M. El-Horbaty, Abdel-Badeeh M. Salem. Classification using deep learning neural networks for brain tumors. Future Computing and Informatics Journal.Volume 3, Issue 1, 2018, Pages 68–71, ISSN 2314-7288.https://doi.org/10.1016/j.fcij.2017.12.001

22. Heba M, El-Dahshan ESA, El-Horbaty ESM, Abdel-Badeeh M (2018) Classification using deep learning neural networks for brain tumors. Future Comput Inf J 3(1):68–71

